# Protein aggregation in wound fluid confines bacterial lipopolysaccharide and reduces inflammation

**DOI:** 10.1101/2023.01.27.525825

**Authors:** Jitka Petrlova, Erik Hartman, Ganna Petruk, Jeremy Chun Hwee Lim, Sunil Shankar Adav, Sven Kjellström, Manoj Puthia, Artur Schmidtchen

**Author notes:** To whom correspondence should be addressed: Jitka Petrlova, Division of Dermatology and Venereology, Department of Clinical Sciences, Lund University, Lund, Sweden. E.H., G.P. and J.C.H.L. contributed equally to this work.

## Abstract

Bacterial lipopolysaccharide (LPS) induces the rapid formation of protein aggregates in human wound fluid. We aimed to define such LPS-induced aggregates and the functional consequences of protein aggregation using a combination of mass spectrometry analyses, biochemical imaging, and experimental animal models. We show that such wound-fluid aggregates contain a multitude of protein classes, including sequences from coagulation factors, annexins, histones, antimicrobial proteins/peptides, and apolipoproteins. Proteins and peptides with a high aggregation propensity were identified, and selected components were verified biochemically by western blot analysis. Staining by thioflavin T and the Amytracker probe demonstrated the presence of amyloid-like aggregates formed after exposure to LPS in vitro in human wound fluid and in vivo in porcine wound models. Using NF-κB-reporter mice and IVIS bioimaging, we show that such wound-fluid LPS aggregates induce a significant reduction in local inflammation compared with LPS in plasma. The results show that protein/peptide aggregation is a mechanism for confining LPS and reducing inflammation and further underscore the connection between host defense and amyloidogenesis.

## Introduction

Skin wounds pose a potential threat for bacterial invasion and sepsis, and a multitude of host defense systems have consequently evolved. These include coagulation, initial hemostasis, and the subsequent action of multiple proteins and peptides of our innate immune system (Esmon *et al*, 2011; Zasloff, 2002). Examples are neutrophil-derived α-defensins and the cathelicidin LL-37 (Frohm *et al*, 1996; Huttner & Bevins, 1999; Zasloff, 2002), histones (Frohm *et al*., 1996; Hirsch, 1958; Hoeksema *et al*, 2016), lysozyme (Frohm *et al*., 1996), and proteolytic products of plasma proteins, such as fibrinogen (Pahlman *et al*, 2013), complement factor C3 (Nordahl *et al*, 2004), and thrombin (Papareddy *et al*, 2010; Saravanan *et al*, 2018).

Lipopolysaccharide (LPS) sensing by Toll-like receptor 4 (TLR4) controls early responses to infection. However, an excessive LPS response is deleterious, causing localized inflammation such as that found in infected wounds, as well as severe systemic responses such as those seen in sepsis (Angus & van der Poll, 2013). Therefore, clearance and control of endotoxins are critical for a robust antibacterial response while maintaining control of inflammatory responses.

We have previously demonstrated that addition of LPS or bacteria to human wound fluids (WFs) leads to precipitation of protein aggregates containing C-terminal thrombin fragments of about 11 kDa, which mediate LPS aggregation and scavenging (Petrlova *et al*, 2017; Petrlova *et al*, 2020). These observations led us to hypothesize that aggregation in wounds constitutes an early sensing and clearance mechanism to control and scavenge excessive LPS levels during wounding and injury. However, it was unknown whether other proteins and peptides apart from the identified thrombin fragments could participate in LPS-scavenging in human WFs.

Thus, we sought to define the LPS interactome in WF and the functional consequences of LPS aggregation. Using mass spectrometry analysis, we show that such aggregates not only contain thrombin sequences, but also other coagulation factors and sequences from protein families, including annexins, histones, antimicrobial proteins/peptides, and apolipoproteins. Proteins and peptides with a high propensity for aggregation were identified, demonstrating a subclass of LPS-interacting molecules in WFs. Selected aggregating components were verified biochemically by western blot analysis. Staining by thioflavin T and the Amytracker probe demonstrates the presence of amyloid-like aggregates formed after exposure to LPS in vitro in human WF, as well as in vivo in porcine wounds. Using NF-κB-reporter mice and IVIS bioimaging, we show that such LPS aggregates in WF induce a significant reduction in local inflammation when compared with LPS administrated with plasma.

## Results

### General experimental outline

The main outline is shown in Figure 1. Briefly, to examine whether LPS induces aggregation and amyloid formation in WFs, we analyzed the supernatant and pellet by biochemical, imaging, and mass spectrometry methodologies. Quantitative analyses were done to measure aggregating proteins, and qualitative studies were done using mass spectrometry analysis of proteomes, western blot of selected proteins, and analyses of the peptidomes generated. Finally, functional analyses were done using in vitro and in vivo analyses of LPS-induced NF-κB activation and the effects of LPS confinement by aggregation.

**Figure 1.**
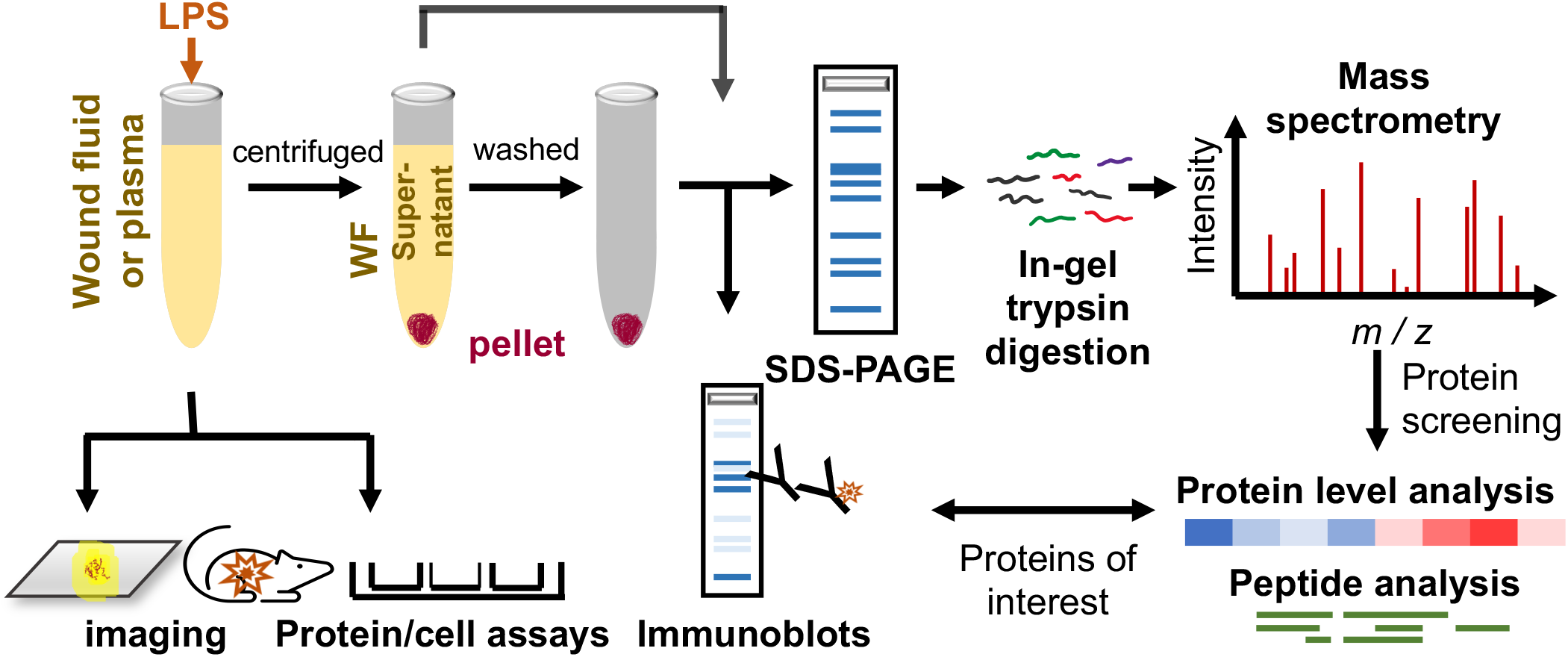
Experimental design. Sample preparation of acute wound fluids, plasma, and pig wound fluid for imaging, analysis, various assays and mass spectrometry.

### LPS induces aggregation in WFs in vitro and in wounds in vivo

Four acute WFs were incubated with LPS, and the protein content in the resulting pellets and supernatants was analyzed. Overall, we detected an increase in protein amounts in pellets of the AWFs after incubation with 100-500 µg/ml of LPS, with three AWFs showing significant changes (Fig 1A, panels on the left). Notably, an increase in the protein amount in the pellet corresponded to a decrease in the supernatants. No such LPS-induced changes were observed in the four plasma samples analyzed (Fig 1A, panels on the right). Moreover, using the fluorescent dye thioflavin T1 (ThT), we detected a significant increase in fluorescence after addition of LPS to the AWFs, but no difference was observed for the plasma samples. ThT specifically binds to β-sheet structures in aggregating proteins and amyloids, so the results demonstrated that LPS caused a significant increase in amyloid-like aggregates in all four AWFs analyzed (Fig 2B).

**Figure 2.**
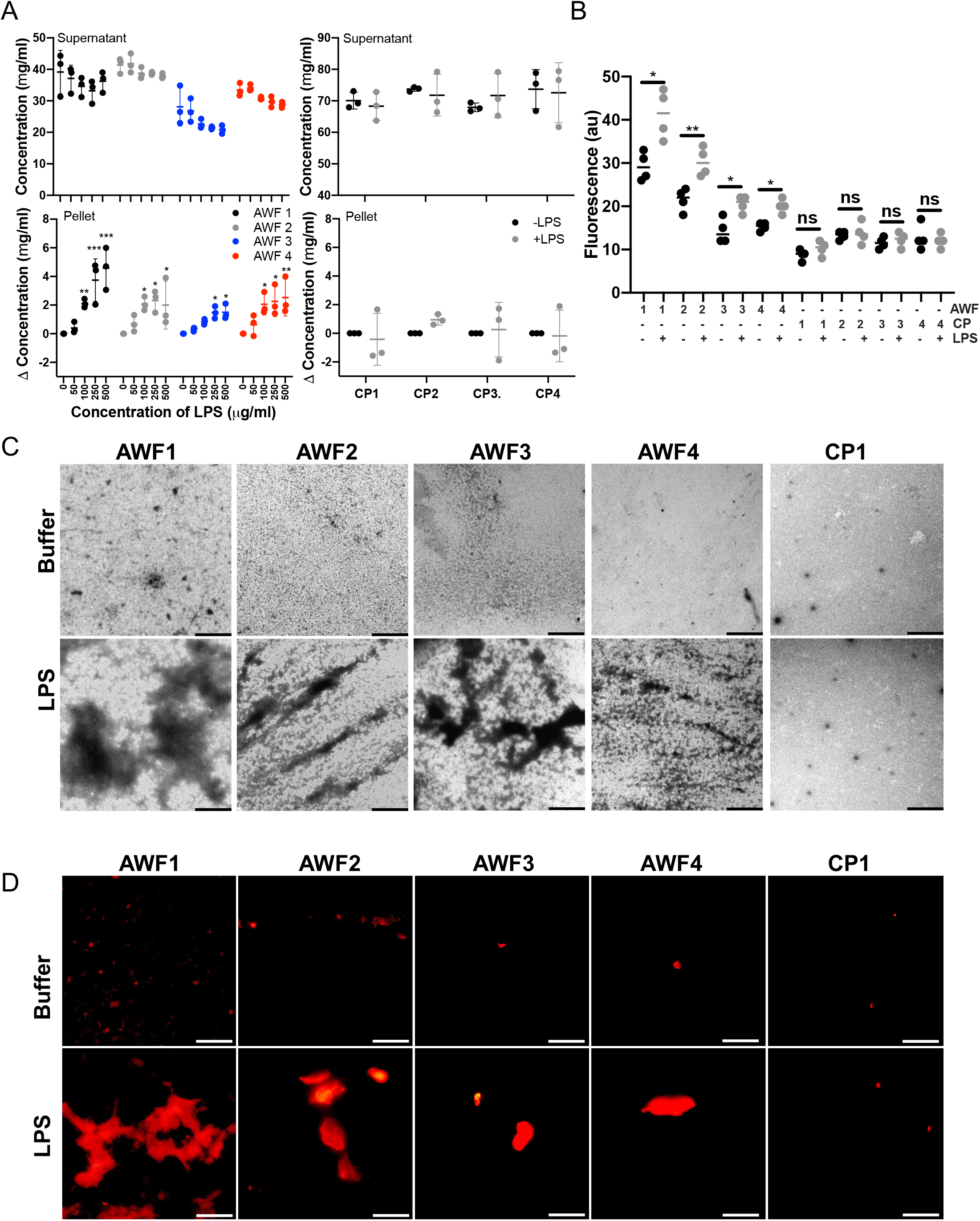
Protein aggregation in AWFs challenged by LPS. A) Analysis of protein content after addition of LPS to acute wound fluids (AWFs). Various doses of LPS were added to AWFs and protein content determined. A significant increase of protein content was detected in the pellets of the AWFs. No increase was observed in the citrated plasma (CP) samples. The graphs are presented as means ± SD from at least three independent experiments. * P ≤ 0.05; **P ≤ 0.01; and ***P ≤ 0.001. P values were determined relative untreated (control) using one-way ANOVA followed by Dunnett’s multiple comparisons test. B) ThT aggregation assay showed a significant increase of aggregates in all AWFs after addition of 100 μg/ml of LPS. No such effects were seen in the CP samples. Statistical analysis was performed using one-way ANOVA with Dunnett’s multiple comparison tests from four independent experiments (n=4). *= P≤ 0.05 and ** = P≤ 0.01. C) TEM – negative stain showed amorphous aggregates in all AWFs exposed to LPS (100 μg/ml). The images represent an example from three independent experiments. The scale bar is 5 μm. D) Fluorescence microscopy using Amytracker 680 staining shows increase of stained aggregates in AWFs after addition of LPS (100 μg/ml). The images represent an example from three independent experiments. The scale bar is 5 μm.

Next, we employed transmission electron microscopy (TEM) with negative stain to visualize the formed protein aggregates, and the results showed that LPS induced formation of aggregates in all AWFs, which were not seen in the CP sample (Fig 1C). Amytracker 680 is a fluorescent tracer for high-quality visualization of protein aggregation and amyloids. Using the fluorescent probe, we detected a significant increase in fluorescence signal in all AWFs subjected to 100 µg/ml of LPS, which again contrasted with the results obtained using CP (Fig 2D and Fig EV1A). Buffer and LPS alone yielded a low fluorescence signal (Fig EV1B).

We next explored whether WF aggregates can also be formed in vivo. For this purpose, we employed a partial thickness wound model in Göttingen minipigs. In a reductionist approach, we used topical LPS application, thus avoiding possible confounders induced by bacterial infection per se. Using Amytracker, we detected a significant increase in aggregates in WFs from LPS-treated wounds in comparison to untreated controls (Figs 3A-B). Likewise, using ThT staining, a higher number of amyloid aggregates was detected in the LPS-treated wounds (Fig 3C). Taken together, the results demonstrate that LPS induces protein aggregation in vitro in human WFs and in vivo in porcine wounds and that the aggregates have amyloid-like properties.

**Figure 3.**
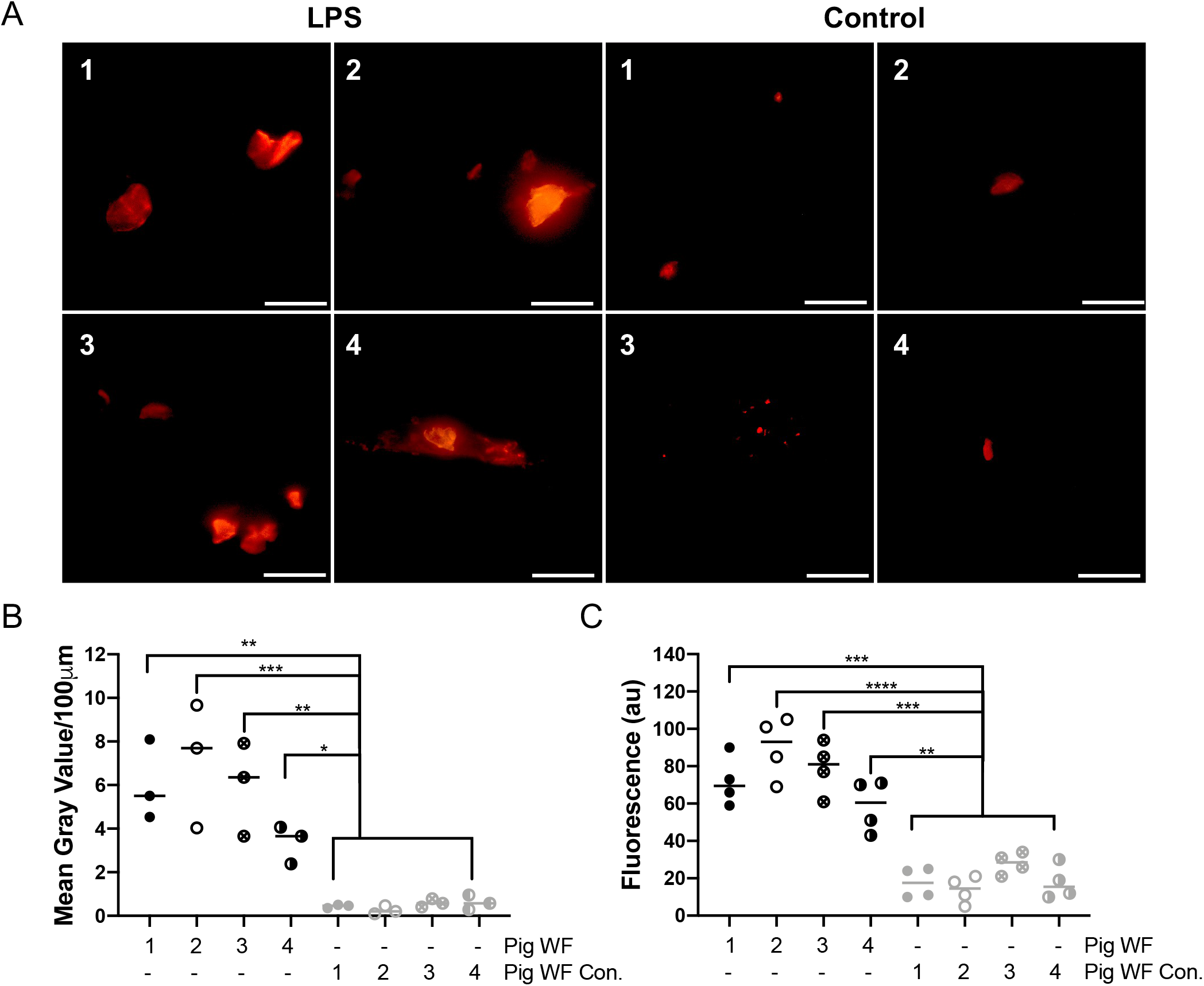
Protein aggregation in porcine wounds in vivo. A) Fluorescence microscopy analysis of samples stained with Amytracker 680 stain shows an increase of aggregates in all AWFs derived from wounds exposed to LPS (100 μg/ml). The images represent an example from three independent experiments. The scale bar is 5 μm. B) Image analyses of Amytracker 680 signal. C) ThT aggregation assay confirmed a significant increase of aggregates in all AWFs from wounds treated with 100 μg/ml of LPS. No aggregates were detected in the control WFs. Statistical analysis was performed using one-way ANOVA with Dunnett’s multiple comparison tests from four independent experiments (n=4). *= P≤ 0.05, ** = P≤ 0.01, *** = P≤ 0.001 and **** = P≤ 0.0001.

### Mass spectrometry analysis of LPS-induced WF aggregates

We next performed a proteomic analysis of the components found in the aggregates with an overall aim of determining whether certain components are prone to aggregation with LPS. For this, protein pellets and supernatants from WFs aggregated with LPS were separated on SDSPAGE and prepared for LC-MS/MS analysis. Peptides from four biological replicates and their technical duplicates were separated and analyzed to determine the specificity of LPS-triggered protein aggregation and enrichment in the pellet.

The mean protein MS intensity in the pellets and supernatants was determined, showing an overall correlation between the intensities of proteins in the pellet and supernatants (Fig 4A). However, the observed spread of the intensities indicated that certain proteins were particularly enriched in the pellet. To visualize this, the relative protein abundance in the pellet was calculated, illustrating the overall aggregation tendency of the individual proteins (Figs 4B and C). We observed that certain sequences derived from annexins, apolipoproteins, hemoglobins, histones, antimicrobial proteins, and coagulation factors were particularly enriched in the LPS-induced WF pellets (Figs 4B and C). The heat maps illustrating all the identified proteins demonstrate the selectivity of aggregation, defining the “LPS aggregatome” (Fig 4D, Fig EV2). To validate the mass spectrometry-based data, we performed SDS-PAGE and western blot analyses. Initial SDS-PAGE analysis showed that the LPS aggregated material in the pellet contained more low-molecular-weight proteins/peptides when compared with the material in the supernatant (Fig 5A). Albumin, a dominating protein in WFs, was observed in the region of 65-70 kDa, and it was noted that its relative abundance was lower in the LPS pellet (Fig 4A), which is in agreement with the mass spectrometry data (Fig EV2). Next, SDS-PAGE followed by western blot analysis demonstrated a relative increase of LL-37, apolipoprotein E (apoE), and histone H2B in the LPS pellet, whereas the opposite was observed for albumin, complement factor C3, and α_1_ antitrypsin. These qualitative data on selected proteins thus supported the mass spectrometry-based results.

**Figure 4.**
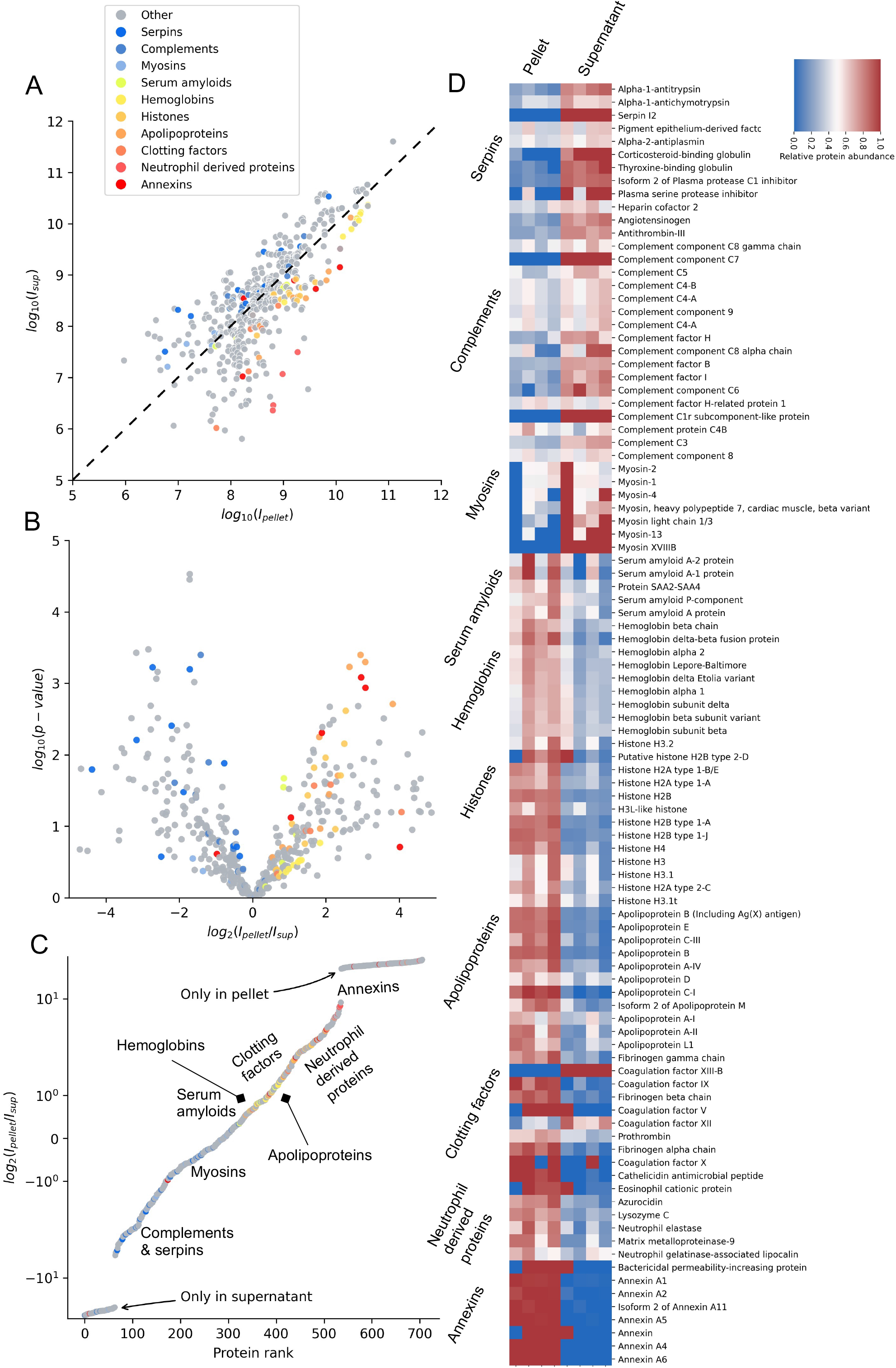
Mass spectrometry proteomic analysis of the LPS aggregatome. A) Protein MS intensities derived from pellets and supernatants of all four AWFs treated with LPS. Each specific protein is represented as a dot in the graph. Proteins which are not at all present in one of the groups are excluded. The dashed diagonal represents equal abundance in both samples. Proteins are colored if they are a part of a relevant protein group. B) A volcano plot of the proteins in the supernatant and pellet. The volcano plot shows the log-transformed p-value over the log-transformed fold change of average protein intensities. Proteins are colored if they are a part of a relevant protein group. C) The log-transformed fold change between pellet and supernatant over the protein rank. Here, the proteins solely present in the pellet and supernatant respectively are included, as contrary to A and B where they are outside the bounds of the axes’ limits. Proteins are colored if they are a part of a relevant protein group. D) Heat map of selected proteins based on the relative protein abundance in the pellet and the supernatant in AWFs subjected to LPS. Proteins were grouped according to protein groups or origins.

**Figure 5.**
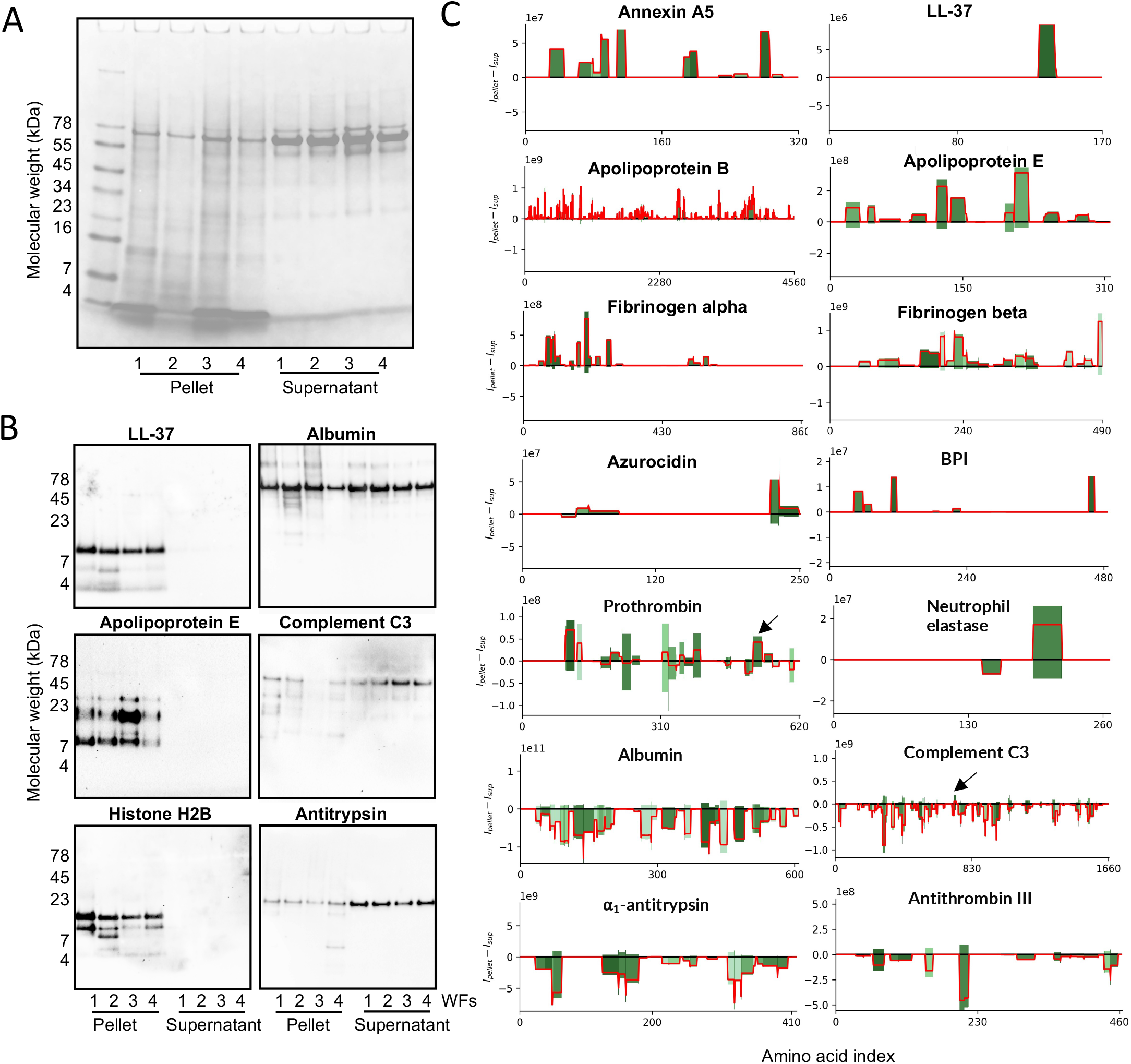
Visualization of proteins and their peptide-fragments. A) SDS-PAGE depicts proteins and peptides in the pellets and supernatants of AWFs subjected to LPS. 30 µg of protein were analyzed. B) Western blot analysis of the selected indicated proteins. The images represent an example from at least three independent experiments. 30 µg of protein were analyzed. C) The x-axis in each protein-plot represents the amino acid index, and the y-axis the summarized intensities in the pellet and supernatant respectively. The intensities in the pellet go in the positive direction (up) and the supernatant the negative (down). The color of the bars corresponds to the number of peptides overlapping each amino acid index, where darker shades of green correspond to a higher overlap. The color cutoffs are defined by quintiles. The red line shows the difference between the intensities in the pellet and supernatant for each amino acid index—i.e., the difference in bar-heights.

Peptide data can be illustrated in peptigrams showing peptide intensity and position in the protein sequence. Therefore, peptigrams of the peptide sequences identified were generated from a set of selected proteins representing various protein classes and grades enrichment in the LPS aggregatome (Fig 5C). We also wanted to probe whether certain regions from these proteins were enriched, so the peptide intensities were subtracted to visualize the differences between the peptidomes of the LPS pellets and the supernatants (Fig 5C, red line). The results clearly demonstrate the selective enrichment and exclusion of peptide sequences, further illustrating the selectivity of LPS-induced aggregation.

### Effects of AWFs on LPS responses in vitro and in vivo

We explored the functional significance of LPS-induced aggregation. For this, we determined the NF-κB/AP-1 activation in THP-1-XBlue-CD14 reporter monocytes. The results showed that all four AWFs inhibited LPS-induced NF-κB activation relative to the buffer and plasma controls. Lepirudin plasma was used to avoid scavenging effects on Ca^2+^ in the cell medium. The AWFs did not affect cell viability (Fig 6A).

**Figure 6.**
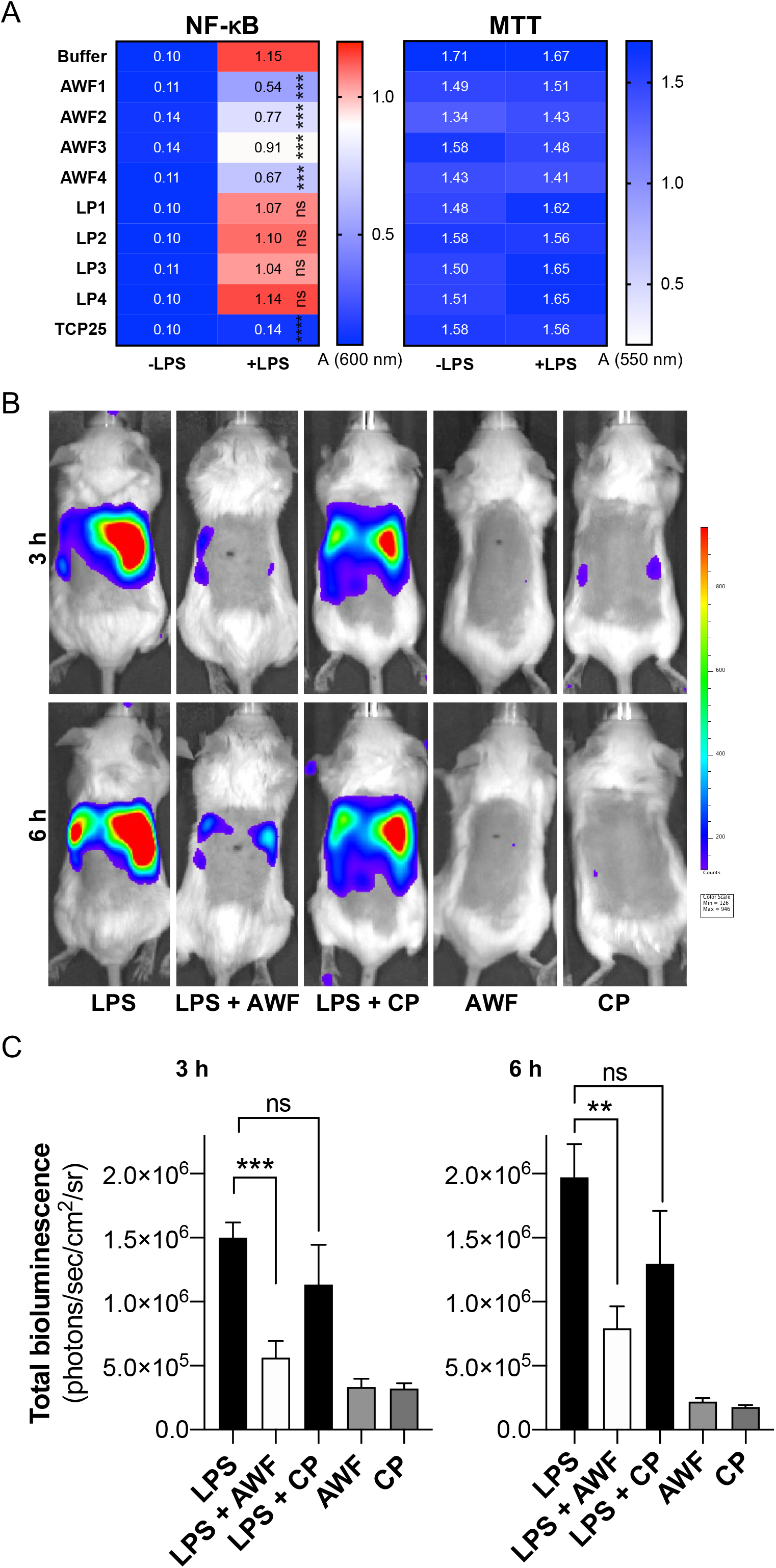
Effects of AWF on LPS-responses in vivo. A) THP-1 cells treated with AWFs and LPS exhibited significantly lower activation of NF-κB compared to LPS alone. No reduction of LPS-induced NF-κB activation is seen after addition of lepirudin plasma (LP). MTT viability assay showed no toxic effect of LPS, AWFs or LPs on THP-1 cells. Statistical analysis was performed using one-way ANOVA with Dunnett’s multiple comparison tests from six independent experiments (n=6). *** = P≤ 0.001, **** = P≤ 0.0001, ns = not significant. E) LPS aggregated by AWF shows reduced NF-κB activation in a mouse model (A). Mice injected subcutaneously with AWF and LPS show significantly lower NF-κB activation when compared with LPS alone or LPS with citrate plasma (CP). All in vivo data are presented as the mean ± SEM (*n* = 5-6 mice). * = P:: 0.05, *P* values were determined using the Mann-Whitney U test.

We next explored whether AWFs could suppress LPS-triggered local inflammation *in vivo*. Using mice with NF-κB activation, we found that AWF was able to significantly reduce LPS-induced inflammation when compared with plasma after 3 and 6 hours (Figs 6B and C). Taken together, these results showed that aggregation of LPS by AWFs reduces NF-κB activation in monocytes in vitro, as well as in experimental mouse models of inflammation.

## Discussion

LPS is a highly proinflammatory substance from Gram-negative bacteria that causes excessive inflammation if not controlled, and we have reported a simple but effective mechanism by which WF can aggregate and reduce LPS effects. Using NF-κB-reporter mice and IVIS bioimaging, we showed that LPS aggregates in WF induce a significantly reduced local inflammation when compared with LPS administrated with plasma. Such aggregates were only observed in porcine wounds challenged with LPS, which underscores the physiological relevance of aggregation in host defense. Many of the proteins and peptides enriched in the LPS pellet (Fig 4, EV Fig 2), such as hCAP18/LL-37, thrombin, other coagulation factors, apoE, apoB, azurocidin, bacterial permeability increasing protein, annexin A5, eosinophil cationic protein, lysozyme, and histones, which are all known for their ability to interact with LPS or LPS-rich bacterial surfaces (Augusto *et al*, 2003; Elsbach & Weiss, 1993; Gaglione *et al*, 2017; Kasetty *et al*, 2011; Lindmark *et al*, 1999; Ohno & Morrison, 1989; Papareddy *et al*., 2010; Petrlova *et al*., 2020; Petruk *et al*, 2021; Pulido *et al*, 2016; Puthia *et al*, 2022; Rand *et al*, 2012; Zasloff, 2002) (see also Table EV1). This indicates a certain degree of LPS selectivity and demonstrates the relevance of the experimental system used.

The aggregates formed after exposure to LPS in vitro in human WF and in vivo in porcine wounds were stained by the amyloid-specific ThT and the Amytracker probe. This demonstrated the amyloid nature of the wound-fluid LPS aggregates. Interestingly, the capacity to aggregate was observed in human WF and not in plasma. As acute WF contains proteases such as neutrophil elastase and various matrix metalloproteinases, it is plausible that proteolytic events may modify certain holoproteins, enabling their LPS interactions and subsequent aggregation. In agreement with this, SDS-PAGE analysis of the LPS aggregatome showed that it contained low molecular fragments that were not observed in the supernatant (Fig 5A). Moreover, this reasoning is elegantly exemplified by prothrombin and thrombin, which yields 11 kDa C-terminal peptides forming aggregates with LPS upon proteolysis (Petrlova *et al*., 2017).

Interestingly, a close inspection of the thrombin peptigram shows that peptides derived from this C-terminal region are particularly enriched (Fig 5C, arrow). Intriguingly, a peptide region from an LPS-interacting part of C3a, VFLDCCNYITELRRQ (Nordahl *et al*., 2004), was also identified in the LPS pellet, whereas the majority of C3 sequences were found in the supernatant (Fig 5C, arrow). However, it cannot be directly assumed that all enriched proteins in the pellet interact with LPS per se, as proteins may co-aggregate in relation to fibrillar amyloids (Bondarev *et al*, 2018; Juhl *et al*, 2019).

Proteomic analyses of plaques from patients with Alzheimer’s disease (AD) (Afagh *et al*, 1996; McGeer *et al*, 1994) and materials from other local and systemic amyloid diseases (Cardoso *et al*, 2002; Karring *et al*, 2013) present lists of co-aggregated proteins comprising many components found here in the LPS aggregatome, including proteases, coagulation factors, and various apolipoproteins. The similarities could indicate a general response of proteins to the formation of fibrillar assemblies (Juhl *et al*., 2019). Therefore, the LPS aggregatome may be viewed as a primordial fast-acting defense system in wounds that is activated by proteolysis and characterized by direct LPS-interactions and likely indirect co-aggregation.

From a pathophysiological perspective, our result on the LPS aggregatome underscores the connection between host defense and amyloidogenesis (Harris *et al*, 2012; Kumar *et al*, 2016; Lee *et al*, 2020; Pasupuleti *et al*, 2009; Soscia *et al*, 2010). Consequently, overactivation of host defense in chronic inflammatory states may induce aberrant aggregation. Therefore, it is notable that neurodegenerative diseases such as Parkinson’s, Huntington’s, and Alzheimer’s disease (AD) have been reported to contain several components of the LPS aggregatome (Table EV1). For example, apolipoprotein E is detected together with amyloid-ß peptides in AD patients (Cedazo-Minguez & Cowburn, 2001).

Correspondingly, peptides from the receptor and lipid-binding regions of apoE (138-150 and 244-272, respectively), from which synthetic peptides have been shown to exhibit antibacterial and immunomodulatory effects (Forbes *et al*, 2013; Wang *et al*, 2013), were found in the LPS aggregatome. Moreover, serum amyloid P, which is enriched in the LPS induced aggregate, has been reported to interact with bacteria (Hind *et al*, 1985) and found in AD and systemic amyloidoses (Tennent *et al*, 1995). Other proteins with well-described roles in host defense— thrombin (Petrlova *et al*., 2017), histones (Hoeksema *et al*., 2016), and LL-37 (Zasloff, 2002)— are also associated with AD or bind to amyloids (De Lorenzi *et al*, 2017; Grammas & Martinez, 2014; Potempska *et al*, 1993) (see also Table EV1).

Evidence shows that dysregulation of the immune system characterizes AD, and the disease could be viewed as a systemic disease that involves activation of peripheral and central immunity. Notably, both acute and chronic systemic inflammation is associated with an increase in cognitive decline in Alzheimer’s disease (Bettcher *et al*, 2021; Holmes *et al*, 2009). Many of the known protein components of AD are present in the LPS aggregatome. This further underscores that fundamental and evolutionarily old host-defense mechanisms designed for protection during injury and wounding can be dysfunctionally activated in other organ systems. This increases the risk for developing neurodegenerative diseases or other systemic and local amyloid diseases. Therefore, studies on the LPS aggregatome may not only yield new pathophysiological insights but could also lead to the discovery of biomarkers for amyloid disease and new strategies to target dysfunctional and ectopic protein aggregation.

## Materials and Methods

### Cell culture and animals

THP-1 cells (ATCC) were cultured in RMPI 1640-GlutaMAX-1 (Gibco, Life Technology ltd., UK). The media was supplemented with 10% (v/v) heat-inactivated FBS (FBSi, Invitrogen, USA) and 1% (v/v) antibiotic-antimycotic solution (AA, Invitrogen) at 37°C in 5% CO_2_. BALB/c tg(NF-κB-RE-Luc)-Xen reporter male mice (Taconic Biosciences) were used for the subcutaneous inflammation experiments and bioimaging. Female Göttingen minipigs (14–16 kg body weight) were used for the partial thickness wound models. The animals were housed under standard conditions of light and temperature and had free access to standard laboratory chow and water.

### Ethics statement

The use of human wound materials and blood was approved by the Ethics Committee at Lund University (LU 708-01 and LU 509-01; permit no. 657-2008). Informed consent was obtained from all donors. All animal experiments were performed according to the Swedish Animal Welfare Act (SFS 1988:534) and were approved by the Animal Ethics Committee of Malmö/Lund, Sweden (permit numbers M88-91/14, M5934-19, M8871-19). Animals were kept under standard conditions of light and temperature with water provided ad libitum.

### Biological materials

Sterile acute wound fluids (AWFs) were obtained from surgical drainages after surgery as described previously (Lundqvist *et al*, 2004). The AWFs were centrifuged, aliquoted, and stored at −20°C. Blood was collected from healthy donors and used for generation of citrated plasma (CP).

### Aggregation of WF proteins by lipopolysaccharide

We incubated four AWFs and four CP samples (100 µl) with increasing concentrations of *Escherichia coli* LPS (0, 50, 100, 200, and 500 μg/ml) (Sigma, USA) in 10 mM Tris for 30 min at pH 7.4 and 37°C. The samples were spun at 10,000 x g for 5 min, and the pellets and supernatants were collected separately. The pellets were washed with 10 mM Tris pH 7.4 buffer and suspended in 100 μl of 8 M urea. Resuspended pellets and 100x diluted supernatants were analyzed for protein concentration using the bicinchoninic acid (BCA) assay according to the manufacturer’s protocol (Thermo Scientific, USA).

For mass spectrometry, AWFs (500 µl) from four different patients with LPS (50 µg) were incubated at 37°C for 30 min. The mixtures were centrifuged at 10000 x g for 5 min, and the pellets and supernatants were collected separately. The pellets were washed with 10 mM Tris pH 7.4 buffer and suspended in a buffer comprising 2% SDS and 50 mM ammonium acetate at pH 6.0. The protein content was determined by the BCA assay.

### Thioflavin T binding assay

The fluorophore thioflavin T (ThT) (Sigma, USA) was obtained from a 1 mM stock stored in the dark at 4°C. We incubated four AWFs and CPs (20 μl) with or without LPS from *E. coli* (100 μg/ml) in 10 mM Tris at pH 7.4 for 30 min at 37°C. We added 180 μl of ThT from the stock solution to a final concentration of 100 μM in 10 mM Tris at pH 7.4 and incubated it for 15 min in the dark. We measured ThT fluorescence using a VICTOR3 multilabel plate counter spectrofluorometer (PerkinElmer, USA) at an excitation of 450 nm with excitation and emission slit widths of 10 nm. The signal obtained from LPS in 10 mM Tris at pH 7.4 was defined as the baseline and subtracted from each AWF and CP sample.

### Transmission electron microscopy – negative stain (TEM)

Aggregates of AWFs were visualized using TEM (Jeol Jem 1230; Jeol, Japan) in combination with negative staining after incubation with LPS (100 μg/ml) or buffer. Images of 10% of AWFs in the presence or absence of LPS (100 μg/ml) were obtained after incubation for 30 min at 37°C. The grids were rendered hydrophilic via glow discharge at low air pressure. Samples were adsorbed onto carbon-coated grids (copper mesh, 400) for 60 s and stained with 7 µl of 2% uranyl acetate for 30 s. For the mounted samples, 10 view fields were examined on the grid (pitch 62 μm) from three independent sample preparations. The CP sample was used in the same experiment as a negative control.

### Minipig partial thickness wound model and WF extraction

We used a minipig partial-thickness wound model to study the LPS-induced aggregation in vivo. Female Göttingen minipigs (14–16 kg body weight) were used. Wounds were induced as described previously (Puthia *et al*, 2020; Stromdahl *et al*, 2021). All procedures were performed by a veterinarian under strict aseptic conditions. The wounding procedures were performed under general anesthesia, and respiratory support was provided with oxygen. Hair on the back was clipped, and the area was thoroughly cleaned with chlorhexidine (MEDI-SCRUB sponge; Rovers, Oss, Netherlands). The back was then shaved and disinfected with chlorhexidine solution (4%) and dried with gauze. Partial-thickness (750-mm deep) wounds (2.5 × 2.5 cm) were made on the back with the help of an electric dermatome (Zimmer).

LPS stock solution was prepared in sterile water (5 mg/ml). A 20-ml amount of stock solution (100 mg LPS) was mixed in 80 ml of a hydrogel composed of 2% hydroxyethyl cellulose (HEC) and applied directly onto the fresh wound. Blank HEC hydrogel (100 ml) was applied to the control wounds. After 15 minutes, the wounds were covered with a primary foam dressing (Mepilex transfer; Mölnlycke Healthcare, Gothenburg, Sweden), followed by a transparent breathable fixation dressing (Mepore film; Mölnlycke, Gothenburg, Sweden). The film was secured using skin staples (Smi, St. Vith, Belgium). For further protection, the wound was then covered with sterile cotton gauze and finally secured with adhesive tape and flexible self-adhesive bandage (Vet Flex, Kruuse, Denmark).

Animals were housed individually and examined daily. Wound dressings were changed daily, and new hydrogel with LPS or control hydrogel was applied to the wounds. Wound dressings were aseptically collected, immediately transferred to a 5-ml pre-chilled tube, and kept on ice. For the extraction of WF, dressings were soaked in 500 ml of ice-cold 10 mM Tris buffer at pH 7.4 and centrifuged for 5 min (2000 x *g*, 4°C). Extracted WFs were aliquoted and stored at −80°C until further analysis.

Polyurethane wound dressings from the porcine partial-thickness wounds were collected aseptically. Dressings were immediately transferred to a 5-ml pre-chilled tube and kept on ice. For the extraction of the porcine wound fluids (PWFs), dressings were soaked in 500 µl of ice-cold 10 mM Tris buffer at pH 7.4 and centrifuged for 5 min (2000 x *g*, 4°C). Extracted PWFs were aliquoted and stored at −80°C until further analysis. Extracted WFs (10 μl) were stained with Amytacker 680 as described in the fluorescence microscopy experiment below.

### Fluorescence microscopy

We used Amytracker 680 (Ebba Biotech, Lund, Sweden) staining to visualize amyloid formation of proteins in WFs. We incubated WFs (10 μl) with LPS (100 μg/ml) for 30 min at 37°C. The samples were subsequently incubated with 50 μl of Amytracker 680 (1000x dilution from the stock solution) in the tube for an additional 30 min of incubation at 37°C. Next, the samples were washed, transferred onto glass slides coated with L-lysine (SIGMA St. Louis, MO, USA), and mounted on microscope slides with fluorescent mounting media (Dako North America, Carpinteria, CA, USA). Ten view fields (1×1 mm) were examined from three independent sample preparations using a Zeiss AxioScope A.1 fluorescence microscope (objectives: Zeiss EC Plan-Neofluar 40×; camera: Zeiss AxioCam MRm; acquisition software: Zeiss Zen 2.6 [blue edition]anti). The CP sample was used in the same experiment as a negative control. The size of aggregates was analyzed as the mean of gray value/μm^2^ ± SEM by ImageJ 1.52k, after all the images were converted to 8-bit and the threshold was adjusted.

### NF-κB activity assay

NF-κB/AP-1 activation in THP-1-XBlue-CD14 reporter monocytes was determined after 20– 24 h of incubation according to the manufacturer’s protocol (InvivoGen, San Diego, USA). Briefly, 1×10^6^ cells/ml in RPMI were seeded in 96-well plates (180 μl) and incubated with AWFs or lepirudin plasma (20 μl) with or without LPS (10 ng/ml) overnight at 37^◦^C in 5% CO_2_ in a total volume of 200 μl. NF-κB activation was determined after 20 h of incubation according to the manufacturer’s instructions by mixing 20 μl of supernatant with 180 μl of SEAP detection reagent (Quanti-BlueTM, InvivoGen, San Diego, CA, USA). The plates were incubated for 2 hours at 37°C, and the absorbance was measured at 600 nm in a VICTOR3 Multilabel Plate Counter spectrofluorometer. Thrombin derived C-terminal peptide (TCP25, Ambiopharm, North Augusta, SC, USA) was used as a positive control for scavenging of LPS-induced NF-κB activation.

### MTT viability assay

A solution sterile filtered solution of MTT (3-(4,5-dimethylthiazolyl)-2,5-diphenyltetrazolium bromide; 5 mg/mL in PBS; Sigma-Aldrich) was stored in the dark at −20°C until usage. We added 20 µl of MTT solution to the remaining overnight culture of THP-1-XBlue-CD14 reporter monocytes from the NF-κB activity assay in 96-well plates, which were incubated at 37°C for 1 hour. The supernatant was then removed, and the blue formazan product generated in the cells was dissolved by the addition of 100 µl of 100% DMSO to each well. The plates were then gently shaken for 10 min at room temperature to dissolve the precipitates. The absorbance was measured at 550 nm in a VICTOR3 Multilabel Plate Counter spectrofluorometer.

### Mouse inflammation model

BALB/c tg(NF-κB-RE-Luc)-Xen reporter mice (Taconic, 10–12-weeks old) were used to study the effects of AWF1 on inflammation (200 μl) after subcutaneous co-treatment with LPS (*E. coli*, 25 μg). The samples were pre-incubated for 30 min at 37°C before injection. The dorsa of the mice were carefully shaved and cleaned. The mice were anesthetized with isoflurane, and 200 μl of the sample was injected subcutaneously. The animals were immediately transferred to individually ventilated cages (IVC) and imaged 3 h later.

We used an In Vivo Imaging System (IVIS) for the determination of NF-κB activation, which plays a key role in the regulation of immune responses during infection. Bioluminescence from the mice was detected and quantified using Living Image 4.0 Software (PerkinElmer). Mice were administrated 100 μl of D-Luciferin intraperitoneally 15 minutes before IVIS imaging (150 mg/kg body weight) (Puthia *et al*., 2020).

### LC-MS/MS analysis

The protein pellets and supernatants (40 µg total protein) from the WFs aggregated with LPS were separated using sodium dodecyl sulfate-polyacrylamide gel electrophoresis (SDS-PAGE; 5% polyacrylamide stacking gel with 12% polyacrylamide separating gel). Electrophoretic separation was performed at 80 V for 20 min and then at 100 V for another 80 min. The gel was then fixed and stained with Coomassie brilliant blue. Next, each sample lane was sliced separately, cut into small pieces of approximately 1 mm^2^, and subjected to complete destaining. After destaining, the gel pieces were reduced with 10 mM DTT, alkylated using 55 mM iodoacetamide, dehydrated with 100% acetonitrile, and then subjected to overnight digestion at 37°C with sequencing-grade modified trypsin (Promega, Madison, WI, USA). The peptides were extracted, vacuum dried, and reconstituted in 0.1% formic acid for LC-MS/MS analysis.

Peptides from four biological replicates and their technical duplicates were separated and analyzed in an LC-MS/MS system comprising of a Dionex Ultimate 3000 RSLC nanoHPLC system, coupled to an online Q-Exactive mass spectrometer (Thermo-Fisher Scientific, USA). Five microliters of each sample were injected into an acclaim peptide trap column via the auto-sampler of the Dionex RSLC nano-HPLC system. The mobile phase A (0.1% FA in 5% acetonitrile) and mobile phase B (0.1% FA in acetonitrile) were used to establish a 60minute gradient, and the flow rate was maintained at 300 nl/ml. Peptides were analyzed using a Dionex EASY-spray column (PepMap® C18, 3um, 100 A) using an EASY nanospray source. The electrospray potential was set at 1.6 kV.

A full MS scan in the range of 350-1600 m/z was acquired at a resolution of 70,000 at m/z 200 with a maximum ion accumulation time of 100 ms. The dynamic exclusion was set to 30 seconds. The resolution for MS/MS spectra was set to 35,000 at m/z 200. The AGC setting was 1E6 for the full MS scan and 2E5 for the MS2 scan. The 10 most intense ions above a count threshold of 1000 were chosen for higher energy collision dissociation (HCD) fragmentation. The maximum ion accumulation time was 120 ms, and an isolation width of 2 Da was used for the MS2 scan. Single and unassigned charged ions were excluded from MS/MS. For HCD, the normalized collision energy was set to 28, and the underfill ratio was defined as 0.1%.

### Mass spectrometry data analysis

Acquired data were processed using Proteome Discoverer (PD) v1.4 (Thermo Scientific, San Jose, CA, USA) with deisotope and deconvolution in MS/MS. The raw files were directly imported into PD and further processed using the designed workflow. Briefly, this workflow includes five processing nodes numbered from 0 to 5. Node 0 is named “spectrum file” and allows selecting raw files, while node 1 is labeled as “spectrum selector” and extracts, deisotopes, and deconvolutes the spectra within a retention time window and precursor ion mass window.

Node 2 was an MS spectrum processor, while node 3 selected the search engine SequestHT, and node 4 used Mascot with database search parameters. The parameter settings were the following: enzyme: trypsin, maximum miss cleavage: 2, minimum peptide length: 6, maximum peptide length: 144, precursor mass tolerance: 10 ppm, fragment mass tolerance:

0.02 Da, dynamic modification: deamidation of Q and N. Node 5 was called “percolator,” and the target FDR (strict) was set as 0.01, while the target FDR (relaxed) was set as 0.05. A database search was performed against the UniProt human database release 2017_02.

For peptidomic analyses, peptide files generated after searching the raw data with PEAKS X were used. For this search, the same modifications and mass tolerances as above were used except that we applied the “no enzyme” option, allowing unspecific cleavages. A database search was performed against the UniProt human database release 2019_01 using 1% FDR and a minimum of two peptides per protein. The obtained peptide/protein lists were exported to Microsoft Excel and analyzed using algorithms in Python 3.9.

The mean protein intensities between the pellets and supernatants collected from all four AWFs were calculated and used to visualize and determine the specificity of LPS-triggered protein aggregation. To visualize the relative discrepancies in protein abundancy between pellet and supernatant, the intensity fraction was defined as follows:

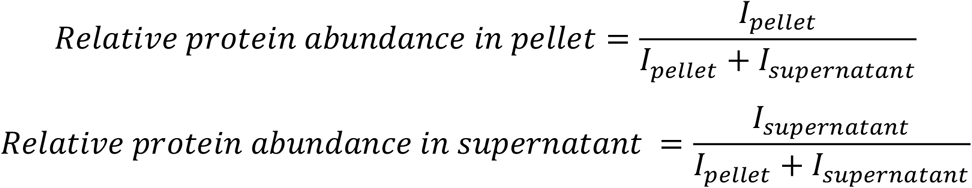

These abundances were calculated for each AWF separately and visualized in heat maps.

The peptidomic content of the pellet and supernatant were visualized using a common convention where the amino acid indexes of proteins are plotted against the abundance of overlapping peptides covering the given index. For a given protein, the abundance *a*_*i*_ of an amino acid at position *i* is the sum of peptide intensities for all peptides that overlap at *i*. The amino acid abundances were then summarized for samples in the pellet and supernatant, resulting in two vectors with the abundance for each amino acid index for each protein.

### SDS-PAGE and western blot

We separated 30 µg of proteins from pellets and supernatants of each AWF using sodium dodecyl sulfate-polyacrylamide gel electrophoresis (Tricine; 10% polyacrylamide stacking gel with 20% polyacrylamide separating gel, Novex by Life Technologies, Carlsbad, CA, USA). Electrophoretic separation was performed at 100 V for 110 min. The gels were fixed and stained with Blue Safe Protein Stain (Thermo Scientific, Rockford, IL, USA) or transferred to a PVDF membrane using a Trans-Blot Turbo system (Bio-Rad, Hercules, CA, USA). The following primary antibodies were used for the western blot experiments: mouse monoclonal antibody against human apolipoprotein E (dilution 1:1000; Abcam, Cambridge, UK), and polyclonal rabbit antibodies against the LL37 epitope (diluted 1:1000; Innovagen AB, Lund, Sweden), human histone 2b (diluted 1:1000; Abcam, Cambridge, UK), human albumin (diluted 1:1000; Dako, Carpinteria, CA, USA), complement C3 (diluted 1:1000; Innovagen AB, Lund, Sweden), and human α_1_ antitrypsin (diluted 1:1000; Dako, Carpinteria, CA, USA). Next, antirabbit or anti-mouse HRP-conjugated antibodies (1:2000; Dako, Carpinteria, CA, USA) were used for detection of the primary antibodies. The proteins and peptides were visualized using chemiluminescent substrates (Thermo Scientific, Rockford, IL, USA) using a ChemiDoc MP imaging system (Bio-Rad, Hercules, CA,).

### Statistical analysis

The graphs of BCA, ThT, and NF-κB activity assays are presented as means ± SD from at least three independent experiments. We assessed differences in these assays using a one-way ANOVA with Dunnett’s multiple comparison tests. Data were analyzed using GraphPad Prism (ααGraphPad Software, Inc., USA) and Python 3.9. P-values less than 0.05 were considered statistically significant (* P<0.05, ** P<0.01, *** P<0.001, and **** P<0.0001).

## Data availability

The mass spectrometry proteomics data have been deposited at the ProteomeXchange Consortium via the PRIDE [1] partner repository with the dataset identifier PXD030521.

Username

## Acknowledgements

This work was supported by grants from the Swedish Research Council (project 2017-02341, 2020-02016), the Leo Foundation, the Welander-Finsen, Crafoord, Österlund, Johanssons, Hedlunds, Kockska, and Söderberg Foundations, the Knut and Alice Wallenberg Foundation, The Swedish Government Funds for Clinical Research (ALF), and the Medical faculty of Lund University. We thank Ann-Charlotte Strömdahl for excellent technical assistance with the THP-1 cell line and Charlotte Welinder for uploading MS data to ProteomeXchange. We acknowledge the Lund University Bioimaging Centrum (LBIC) for access to electron microscopy facilities. The mass spectrometry proteomics data have been deposited at the ProteomeXchange Consortium via the PRIDE [1] partner repository with the dataset identifier PXD030521.

## Author Contributions

J.P. and A.S. conceived the project and designed the experiments. J.P. performed the experiments for protein detection, aggregation assay, cell assay, and microscopy, including sample preparation for MS experiments and in vivo experiments. G.P., J.C.H.L., and E.H. performed the bioinformatic analysis of MS data, M.P. performed the in vivo experiments, and S.S. carried out the MS experiments. S.K. supervised the bioinformatic analysis of MS data. J.P. and A.S. wrote the manuscript. All the authors discussed the results and commented on the final manuscript.

