## Supplementary information for "Protein aggregation in wound fluid confines bacterial lipopolysaccharide and reduces inflammation"

**Expanded View**

**Table EV1**

Proteins aggregating in the presence of LPS and potential links to LPS binding, antimicrobial activity, amyloid formation, or Alzheimer's disease (AD). Proteins/peptides are shown in falling aggregation index order. Note that the list shows associations, and certain components, such as apolipoproteins or serum amyloids, may be involved in pathogenesis or, in the case of lysozyme, have possible protective effects on amyloid formation or AD development.

| **Protein** | **LPS-interaction** | **Antimicrobial** | **Amyloid** | **AD** |
|---|---|---|---|---|
| Bactericidal/permeability increasing protein | (1,2) | (1) | | |
| Annexin A5 | (3) | | (4) | (4,5) |
| CAMP (LL-37) | (6-8) | (8) | (9,10) | (11,12) |
| Eosinophil cationic protein | (13) | (14) | (14,15) | (16) |
| Fibrinogen alpha | | | (17) | (18,19) |
| Fibrinogen beta | | (20) | | (19) |
| Apolipoprotein B | (21) | (22) | (23) | (24) |
| Apolipoprotein E | (25,26) | (27-29) | (23,30,31) | (32-34) |
| Histone H2B | (35) | (36) | (37) | (38,39) |
| Histone H4 | (35) | (40) | (41) | (39) |
| Azurocidin | (42) | (43,44) | | (45) |
| Lysozyme C | (46) | | (17,47,48) | (49) |
| Neutrophil elastase | | | (50) | (12) |
| Hemoglobin alpha and beta chain | (51) | (52-54) | (55,56) | (57-60) |
| Thrombin | (61) | (62,63) | (64,65) | (66-68) |
| Serum amyloid protein | (69,70) | (70,71) | (72-74) | (72,73,75) |
| NGAL | (76) | (76) | (77) | (78) |
| Histone H3 | (36) | (35,79) | (41) | (80) |
| Apolipoprotein A1 | (81,82) | (83) | (84-86) | (87-89) |

16. Navarro, S., Boix, E., Cuchillo, C. M., and Nogues, M. V. (2010) Eosinophil-induced neurotoxicity: the role of eosinophil cationic protein/RNase 3. *J Neuroimmunol* **227**, 60-70
17. Biza, K. V., Nastou, K. C., Tsiolaki, P. L., Mastrokalou, C. V., Hamodrakas, S. J., and Iconomidou, V. A. (2017) The amyloid interactome: Exploring protein aggregation. *PLoS One* **12**, e0173163
18. Benson, M. D., Liepnieks, J., Uemichi, T., Wheeler, G., and Correa, R. (1993) Hereditary renal amyloidosis associated with a mutant fibrinogen alpha-chain. *Nat Genet* **3**, 252-255
19. Kiddle, S. J., Thambisetty, M., Simmons, A., Riddoch-Contreras, J., Hye, A., Westman, E., Pike, I., Ward, M., Johnston, C., Lupton, M. K., Lunnon, K., Soininen, H., Kloszewska, I., Tsolaki, M., Vellas, B., Mecocci, P., Lovestone, S., Newhouse, S., Dobson, R., and Alzheimers Disease Neuroimaging, I. (2012) Plasma based markers of [11C] PiB-PET brain amyloid burden. *PLoS One* **7**, e44260
20. Pahlman, L. I., Morgelin, M., Kasetty, G., Olin, A. I., Schmidtchen, A., and Herwald, H. (2013) Antimicrobial activity of fibrinogen and fibrinogen-derived peptides--a novel link between coagulation and innate immunity. *Thromb Haemost* **109**, 930-939
21. Vreugdenhil, A. C., Snoek, A. M., van 't Veer, C., Greve, J. W., and Buurman, W. A. (2001) LPS-binding protein circulates in association with apoB-containing lipoproteins and enhances endotoxin-LDL/VLDL interaction. *J Clin Invest* **107**, 225-234
22. Gaglione, R., Cesaro, A., Dell'Olmo, E., Della Ventura, B., Casillo, A., Di Girolamo, R., Velotta, R., Notomista, E., Veldhuizen, E. J. A., Corsaro, M. M., De Rosa, C., and Arciello, A. (2019) Effects of human antimicrobial cryptides identified in apolipoprotein B depend on specific features of bacterial strains. *Sci Rep* **9**, 6728
23. Mullins, R. F., Russell, S. R., Anderson, D. H., and Hageman, G. S. (2000) Drusen associated with aging and age-related macular degeneration contain proteins common to extracellular deposits associated with atherosclerosis, elastosis, amyloidosis, and dense deposit disease. *Faseb J* **14**, 835-846
24. Namba, Y., Tsuchiya, H., and Ikeda, K. (1992) Apolipoprotein B immunoreactivity in senile plaque and vascular amyloids and neurofibrillary tangles in the brains of patients with Alzheimer's disease. *Neurosci Lett* **134**, 264-266
25. Petruk, G., Elven, M., Hartman, E., Davoudi, M., Schmidtchen, A., Puthia, M., and Petrlova, J. (2021) The role of full-length apolipoprotein E in clearance of Gram-negative bacteria and their endotoxins. *J Lipid Res*, 100086
26. Puthia, M., Marzinek, J. K., Petruk, G., Erturk Bergdahl, G., Bond, P. J., and Petrlova, J. (2022) Antibacterial and Anti-Inflammatory Effects of Apolipoprotein E. *Biomedicines* **10**
27. Wang, C. Q., Yang, C. S., Yang, Y., Pan, F., He, L. Y., and Wang, A. M. (2013) An apolipoprotein E mimetic peptide with activities against multidrug-resistant bacteria and immunomodulatory effects. *J Pept Sci* **19**, 745-750
28. Azuma, M., Kojimab, T., Yokoyama, I., Tajiri, H., Yoshikawa, K., Saga, S., and Del Carpio, C. A. (2000) A synthetic peptide of human apoprotein E with antibacterial activity. *Peptides* **21**, 327-330
29. Zanfardino, A., Bosso, A., Gallo, G., Pistorio, V., Di Napoli, M., Gaglione, R., Dell'Olmo, E., Varcamonti, M., Notomista, E., Arciello, A., and Pizzo, E. (2018) Human apolipoprotein E as a reservoir of cryptic bioactive peptides: The case of ApoE 133-167. *J Pept Sci* **24**, e3095

30. Furumoto, H., Hashimoto, Y., Muto, M., Shimizu, T., and Nakamura, K. (2002) Apolipoprotein E4 is associated with primary localized cutaneous amyloidosis. *J Invest Dermatol* **119**, 532-533
31. Furumoto, H., Shimizu, T., Asagami, C., Muto, M., Takahashi, M., Hoshii, Y., Ishihara, T., and Nakamura, K. (1998) Apolipoprotein E is present in primary localized cutaneous amyloidosis. *J Invest Dermatol* **111**, 417-421
32. Cedazo-Minguez, A., and Cowburn, R. F. (2001) Apolipoprotein E: a major piece in the Alzheimer's disease puzzle. *J Cell Mol Med* **5**, 254-266
33. Yamazaki, Y., Zhao, N., Caulfield, T. R., Liu, C. C., and Bu, G. (2019) Apolipoprotein E and Alzheimer disease: pathobiology and targeting strategies. *Nat Rev Neurol* **15**, 501-518
34. Husain, M. A., Laurent, B., and Plourde, M. (2021) APOE and Alzheimer's Disease: From Lipid Transport to Physiopathology and Therapeutics. *Front Neurosci* **15**, 630502
35. Morita, S., Tagai, C., Shiraishi, T., Miyaji, K., and Iwamuro, S. (2013) Differential mode of antimicrobial actions of arginine-rich and lysine-rich histones against Gram-positive Staphylococcus aureus. *Peptides* **48**, 75-82
36. Kawasaki, H., and Iwamuro, S. (2008) Potential roles of histones in host defense as antimicrobial agents. *Infect Disord Drug Targets* **8**, 195-205
37. Du Clos, T. W. (1996) The interaction of C-reactive protein and serum amyloid P component with nuclear antigens. *Mol Biol Rep* **23**, 253-260
38. Zafar, S., Shafiq, M., Younas, N., Schmitz, M., Ferrer, I., and Zerr, I. (2017) Prion Protein Interactome: Identifying Novel Targets in Slowly and Rapidly Progressive Forms of Alzheimer's Disease. *J Alzheimers Dis* **59**, 265-275
39. Lu, X., Wang, L., Yu, C., Yu, D., and Yu, G. (2015) Histone Acetylation Modifiers in the Pathogenesis of Alzheimer's Disease. *Front Cell Neurosci* **9**, 226
40. Lee, D. Y., Huang, C. M., Nakatsuji, T., Thiboutot, D., Kang, S. A., Monestier, M., and Gallo, R. L. (2009) Histone H4 is a major component of the antimicrobial action of human sebocytes. *J Invest Dermatol* **129**, 2489-2496
41. Munishkina, L. A., Fink, A. L., and Uversky, V. N. (2004) Conformational prerequisites for formation of amyloid fibrils from histones. *J Mol Biol* **342**, 1305-1324
42. Heinzelmann, M., Mercer-Jones, M. A., Flodgaard, H., and Miller, F. N. (1998) Heparin-binding protein (CAP37) is internalized in monocytes and increases LPS-induced monocyte activation. *Journal of Immunology* **160**, 5530-5536
43. Shafer, W. M., Martin, L. E., and Spitznagel, J. K. (1984) Cationic antimicrobial proteins isolated from human neutrophil granulocytes in the presence of diisopropyl fluorophosphate. *Infect Immun* **45**, 29-35
44. Pereira, H. A., Erdem, I., Pohl, J., and Spitznagel, J. K. (1993) Synthetic bactericidal peptide based on CAP37: a 37-kDa human neutrophil granule-associated cationic antimicrobial protein chemotactic for monocytes. *Proc Natl Acad Sci U S A* **90**, 4733-4737
45. Pereira, H. A., Kumar, P., and Grammas, P. (1996) Expression of CAP37, a novel inflammatory mediator, in Alzheimer's disease. *Neurobiol Aging* **17**, 753-759
46. Ohno, N., and Morrison, D. C. (1989) Lipopolysaccharide interactions with lysozyme differentially affect lipopolysaccharide immunostimulatory activity. *Eur J Biochem* **186**, 629-636
47. Booth, D. R., Sunde, M., Bellotti, V., Robinson, C. V., Hutchinson, W. L., Fraser, P. E., Hawkins, P. N., Dobson, C. M., Radford, S. E., Blake, C. C., and Pepys, M. B.

(1997) Instability, unfolding and aggregation of human lysozyme variants underlying amyloid fibrillogenesis. *Nature* **385**, 787-793
48. Helmfors, L., Boman, A., Civitelli, L., Nath, S., Sandin, L., Janefjord, C., McCann, H., Zetterberg, H., Blennow, K., Halliday, G., Brorsson, A. C., and Kagedal, K. (2015) Protective properties of lysozyme on beta-amyloid pathology: implications for Alzheimer disease. *Neurobiol Dis* **83**, 122-133
49. Sandin, L., Bergkvist, L., Nath, S., Kielkopf, C., Janefjord, C., Helmfors, L., Zetterberg, H., Blennow, K., Li, H., Nilsberth, C., Garner, B., Brorsson, A. C., and Kagedal, K. (2016) Beneficial effects of increased lysozyme levels in Alzheimer's disease modelled in Drosophila melanogaster. *FEBS J* **283**, 3508-3522
50. Stone, P. J., Campistol, J. M., Abraham, C. R., Rodgers, O., Shirahama, T., and Skinner, M. (1993) Neutrophil proteases associated with amyloid fibrils. *Biochem Biophys Res Commun* **197**, 130-136
51. Bahl, N., Du, R., Winarsih, I., Ho, B., Tucker-Kellogg, L., Tidor, B., and Ding, J. L. (2011) Delineation of lipopolysaccharide (LPS)-binding sites on hemoglobin: from in silico predictions to biophysical characterization. *J Biol Chem* **286**, 37793-37803
52. Parish, C. A., Jiang, H., Tokiwa, Y., Berova, N., Nakanishi, K., McCabe, D., Zuckerman, W., Xia, M. M., and Gabay, J. E. (2001) Broad-spectrum antimicrobial activity of hemoglobin. *Bioorg Med Chem* **9**, 377-382
53. Sheshadri, P., and Abraham, J. (2012) Antimicrobial properties of hemoglobin. *Immunopharmacol Immunotoxicol* **34**, 896-900
54. Deng, L. X., Pan, X. L., Wang, Y., Wang, L. L., Zhou, X. E., Li, M., Feng, Y., Wu, Q., Wang, B. Y., and Huang, N. (2009) Hemoglobin and its derived peptides may play a role in the antibacterial mechanism of the vagina. *Hum Reprod* **24**, 211-218
55. Iram, A., and Naeem, A. (2013) Detection and analysis of protofibrils and fibrils of hemoglobin: implications for the pathogenesis and cure of heme loss related maladies. *Arch Biochem Biophys* **533**, 69-78
56. Heuschkel, M. A., Skenteris, N. T., Hutcheson, J. D., van der Valk, D. D., Bremer, J., Goody, P., Hjortnaes, J., Jansen, F., Bouten, C. V. C., van den Bogaerdt, A., Matic, L., Marx, N., and Goettsch, C. (2020) Integrative Multi-Omics Analysis in Calcific Aortic Valve Disease Reveals a Link to the Formation of Amyloid-Like Deposits. *Cells* **9**
57. Arioz, B. I., Tufekci, K. U., Olcum, M., Durur, D. Y., Akarlar, B. A., Ozlu, N., Bagriyanik, H. A., Keskinoglu, P., Yener, G., and Genc, S. (2021) Proteome profiling of neuron-derived exosomes in Alzheimer's disease reveals hemoglobin as a potential biomarker. *Neurosci Lett* **755**, 135914
58. Kim, J. W., Byun, M. S., Yi, D., Lee, J. H., Jeon, S. Y., Ko, K., Joung, H., Jung, G., Lee, J. Y., Sohn, C. H., Lee, Y. S., Kim, Y. K., and Lee, D. Y. (2021) Blood Hemoglobin, in-vivo Alzheimer Pathologies, and Cognitive Impairment: A Cross-Sectional Study. *Front Aging Neurosci* **13**, 625511
59. Yoo, S. H., Woo, S. W., Shin, M. J., Yoon, J. A., Shin, Y. I., and Hong, K. S. (2020) Diagnosis of Mild Cognitive Impairment Using Cognitive Tasks: A Functional Near-Infrared Spectroscopy Study. *Curr Alzheimer Res* **17**, 1145-1160
60. Gattas, B. S., Ibetoh, C. N., Stratulat, E., Liu, F., Wuni, G. Y., Bahuva, R., Shafiq, M. A., and Gordon, D. K. (2020) The Impact of Low Hemoglobin Levels on Cognitive Brain Functions. *Cureus* **12**, e11378
61. Kalle, M., Papareddy, P., Kasetty, G., Morgelin, M., van der Plas, M. J., Rydengard, V., Malmsten, M., Albiger, B., and Schmidtchen, A. (2012) Host defense peptides of thrombin modulate inflammation and coagulation in endotoxin-mediated shock and Pseudomonas aeruginosa sepsis. *PLoS One* **7**, e51313

62. Tang, Y. Q., Yeaman, M. R., and Selsted, M. E. (2002) Antimicrobial peptides from human platelets. *Infect Immun* **70**, 6524-6533
63. Papareddy, P., Rydengard, V., Pasupuleti, M., Walse, B., Morgelin, M., Chalupka, A., Malmsten, M., and Schmidtchen, A. (2010) Proteolysis of human thrombin generates novel host defense peptides. *PLoS pathogens* **6**, e1000857
64. Petrlova, J., Hansen, F. C., van der Plas, M. J. A., Huber, R. G., Morgelin, M., Malmsten, M., Bond, P. J., and Schmidtchen, A. (2017) Aggregation of thrombin-derived C-terminal fragments as a previously undisclosed host defense mechanism. *Proc Natl Acad Sci U S A* **114**, E4213-E4222
65. Gastineau, D. A., Gertz, M. A., Daniels, T. M., Kyle, R. A., and Bowie, E. J. (1991) Inhibitor of the thrombin time in systemic amyloidosis: a common coagulation abnormality. *Blood* **77**, 2637-2640
66. Akiyama, H., Ikeda, K., Kondo, H., and McGeer, P. L. (1992) Thrombin accumulation in brains of patients with Alzheimer's disease. *Neurosci Lett* **146**, 152-154
67. Arai, T., Miklossy, J., Klegeris, A., Guo, J. P., and McGeer, P. L. (2006) Thrombin and prothrombin are expressed by neurons and glial cells and accumulate in neurofibrillary tangles in Alzheimer disease brain. *J Neuropathol Exp Neurol* **65**, 19-25
68. Zamolodchikov, D., Renne, T., and Strickland, S. (2016) The Alzheimer's disease peptide beta-amyloid promotes thrombin generation through activation of coagulation factor XII. *Journal of Thrombosis and Haemostasis* **14**, 995-1007
69. de Haas, C. J., van Leeuwen, E. M., van Bommel, T., Verhoef, J., van Kessel, K. P., and van Strijp, J. A. (2000) Serum amyloid P component bound to gram-negative bacteria prevents lipopolysaccharide-mediated classical pathway complement activation. *Infect Immun* **68**, 1753-1759
70. Noursadeghi, M., Bickerstaff, M. C., Gallimore, J. R., Herbert, J., Cohen, J., and Pepys, M. B. (2000) Role of serum amyloid P component in bacterial infection: protection of the host or protection of the pathogen. *Proc Natl Acad Sci U S A* **97**, 14584-14589
71. Hind, C. R., Collins, P. M., Baltz, M. L., and Pepys, M. B. (1985) Human serum amyloid P component, a circulating lectin with specificity for the cyclic 4,6-pyruvate acetal of galactose. Interactions with various bacteria. *Biochem J* **225**, 107-111
72. Tennent, G. A., Lovat, L. B., and Pepys, M. B. (1995) Serum amyloid P component prevents proteolysis of the amyloid fibrils of Alzheimer disease and systemic amyloidosis. *Proc Natl Acad Sci U S A* **92**, 4299-4303
73. Coria, F., Castano, E., Prelli, F., Larrondo-Lillo, M., van Duinen, S., Shelanski, M. L., and Frangione, B. (1988) Isolation and characterization of amyloid P component from Alzheimer's disease and other types of cerebral amyloidosis. *Lab Invest* **58**, 454-458
74. Botto, M., Hawkins, P. N., Bickerstaff, M. C. M., Herbert, J., Bygrave, A. E., McBride, A., Hutchinson, W. L., Tennent, G. A., Walport, M. J., and Pepys, M. B. (1997) Amyloid deposition is delayed in mice with targeted deletion of the serum amyloid P component gene. *Nature Medicine* **3**, 855-859
75. Duong, T., Pommier, E. C., and Scheibel, A. B. (1989) Immunodetection of the amyloid P component in Alzheimer's disease. *Acta Neuropathol* **78**, 429-437
76. Flo, T. H., Smith, K. D., Sato, S., Rodriguez, D. J., Holmes, M. A., Strong, R. K., Akira, S., and Aderem, A. (2004) Lipocalin 2 mediates an innate immune response to bacterial infection by sequestrating iron. *Nature* **432**, 917-921
77. Sousa, M. M., do Amaral, J. B., Guimaraes, A., and Saraiva, M. J. (2005) Up-regulation of the extracellular matrix remodeling genes, biglycan, neutrophil

gelatinase-associated lipocalin, and matrix metalloproteinase-9 in familial amyloid polyneuropathy. *Faseb J* **19**, 124-126
78. Naude, P. J., Dekker, A. D., Coppus, A. M., Vermeiren, Y., Eisel, U. L., van Duijn, C. M., Van Dam, D., and De Deyn, P. P. (2015) Serum NGAL is Associated with Distinct Plasma Amyloid-beta Peptides According to the Clinical Diagnosis of Dementia in Down Syndrome. *J Alzheimers Dis* **45**, 733-743
79. Hoeksema, M., van Eijk, M., Haagsman, H. P., and Hartshorn, K. L. (2016) Histones as mediators of host defense, inflammation and thrombosis. *Future Microbiol* **11**, 441-453
80. Narayan, P. J., Lill, C., Faull, R., Curtis, M. A., and Dragunow, M. (2015) Increased acetyl and total histone levels in post-mortem Alzheimer's disease brain. *Neurobiol Dis* **74**, 281-294
81. Biedzka-Sarek, M., Metso, J., Kateifides, A., Meri, T., Jokiranta, T. S., Muszynski, A., Radziejewska-Lebrecht, J., Zannis, V., Skurnik, M., and Jauhiainen, M. (2011) Apolipoprotein A-I exerts bactericidal activity against Yersinia enterocolitica serotype O:3. *J Biol Chem* **286**, 38211-38219
82. Gupta, H., Dai, L., Datta, G., Garber, D. W., Grenett, H., Li, Y., Mishra, V., Palgunachari, M. N., Handattu, S., Gianturco, S. H., Bradley, W. A., Anantharamaiah, G. M., and White, C. R. (2005) Inhibition of lipopolysaccharide-induced inflammatory responses by an apolipoprotein AI mimetic peptide. *Circ Res* **97**, 236-243
83. Tada, N., Sakamoto, T., Kagami, A., Mochizuki, K., and Kurosaka, K. (1993) Antimicrobial activity of lipoprotein particles containing apolipoprotein Al. *Mol Cell Biochem* **119**, 171-178
84. Obici, L., Franceschini, G., Calabresi, L., Giorgetti, S., Stoppini, M., Merlini, G., and Bellotti, V. (2006) Structure, function and amyloidogenic propensity of apolipoprotein A-I. *Amyloid* **13**, 191-205
85. Westermark, P., Mucchiano, G., Marthin, T., Johnson, K. H., and Sletten, K. (1995) Apolipoprotein A1-derived amyloid in human aortic atherosclerotic plaques. *Am J Pathol* **147**, 1186-1192
86. Nichols, W. C., Dwulet, F. E., Liepnieks, J., and Benson, M. D. (1988) Variant Apolipoprotein-a-I as a Major Constituent of a Human Hereditary Amyloid. *Biochem Bioph Res Co* **156**, 762-768
87. Merched, A., Xia, Y., Visvikis, S., Serot, J. M., and Siest, G. (2000) Decreased high-density lipoprotein cholesterol and serum apolipoprotein AI concentrations are highly correlated with the severity of Alzheimer's disease. *Neurobiol Aging* **21**, 27-30
88. Lewis, T. L., Cao, D., Lu, H., Mans, R. A., Su, Y. R., Jungbauer, L., Linton, M. F., Fazio, S., LaDu, M. J., and Li, L. (2010) Overexpression of human apolipoprotein A-I preserves cognitive function and attenuates neuroinflammation and cerebral amyloid angiopathy in a mouse model of Alzheimer disease. *J Biol Chem* **285**, 36958-36968
89. Endres, K. (2021) Apolipoprotein A1, the neglected relative of Apolipoprotein E and its potential role in Alzheimer's disease. *Neural Regen Res* **16**, 2141-2148

**Figure EV1**

A) Image analyses of Amytracker 680 signal in AWFs. B) Fluorescence microscopy analysis using Amytracker 680 stain. No protein aggregation in buffer or LPS alone is detected. The scale bar is 5 µm.

**Figure EV2**

A clustered* heatmap of the proteomic content of the pellet and supernatant. The color represents the relative intensity, i.e., $(I_p/I_{total})$, where $I_p$ is the protein intensity in the pellet or supernatant respectively, and $I_{total}$ is the total intensity in both the pellet and supernatant for the given protein. The map was clustered on both samples and proteins. Proteins not found in 2 samples were discarded (evaluated for the pellet and supernatant individually).

*Clustering was performed using the average Euclidian distance on the relative protein abundances.
