## Supplementary figures and images for "Protein aggregation in wound fluid confines bacterial lipopolysaccharide and reduces inflammation"

### Supplementary information figures

EV figure 1

A

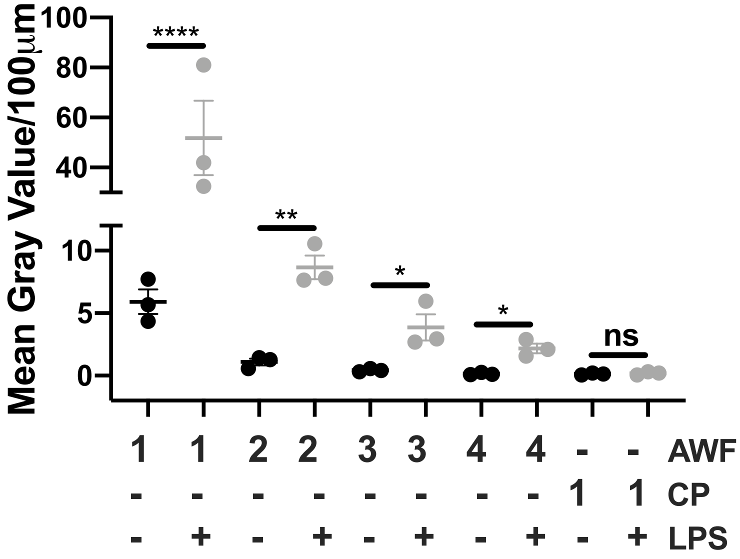


B

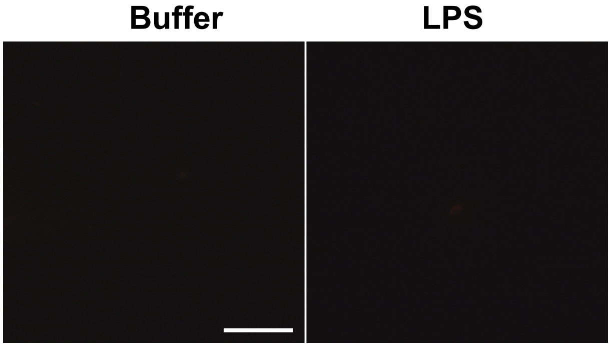

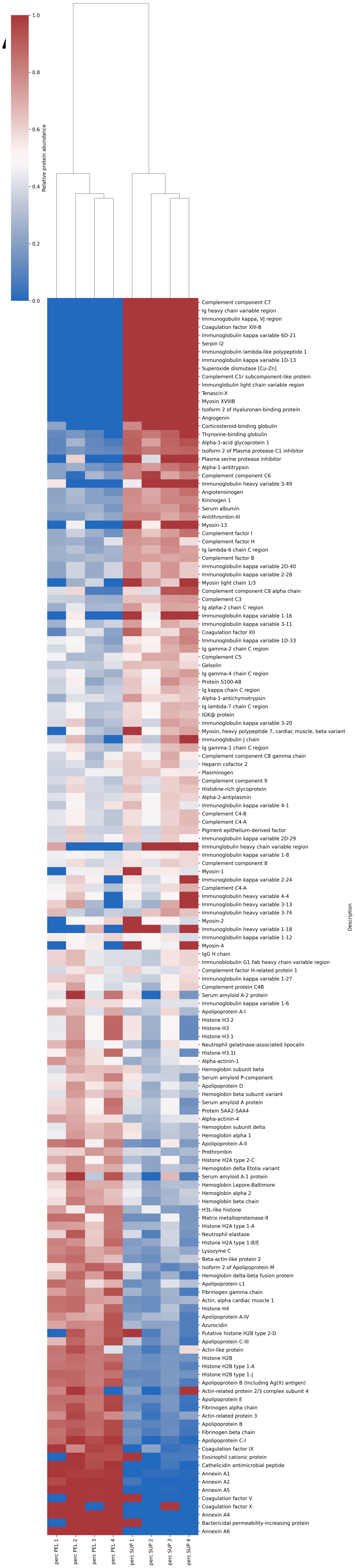
